# Cellular and Network Effects of Introducing Connexin-36 Expression in an Uncoupled Neuronal Population in the Mouse Hypothalamus

**DOI:** 10.64898/2026.09.23.753757

**Authors:** Andriana Mantzafou, Andrea Locarno, Christian Broberger

## Abstract

Electrical synapses are prevalent throughout nervous systems, including the mammalian brain. Several lines of evidence implicate these gap junction connections in several important roles in neural networks. Yet, progress has been hampered by a shortage of experimental tools to investigate their function with sufficient precision. In an effort to address this deficit, we here take advantage of a curious species-difference in electrical coupling in tuberoinfundibular dopamine (TIDA) neurons of the rodent hypothalamus. In rats, these cells exhibit strong electrical coupling, generating stereotyped, slow oscillations that are synchronized across the population. In contrast, mouse TIDA neurons lack gap junctions and display diverse, faster, and asynchronous oscillations. Motivated by this natural discrepancy, we designed a gain-of-function strategy to induce localized and robust expression of the main pore-forming protein of neuronal gap junctions, connexin-36 (Cx36), in mouse TIDA neurons by viral vectors. This perturbation yielded electrical coupling that was weaker than the rat TIDA system, but on par with many other brain populations connected by gap junctions. Cx36 overexpression resulted in significantly increased auto- and cross-correlation, as well as augmented functional connectivity, in agreement with the induction of electrical synapses in the population. Yet, these measures were modest in comparison with rat TIDA neurons, and Cx36 overexpression failed to alter serum levels of the hormone prolactin, which is controlled by this neuroendocrine system. These findings highlight the potential and limitations of experimentally inducing electrical synapses in a natively uncoupled system, and suggest directions for future improvements.

**Significance statement:** Many neurons communicate by electrical synapses, in addition to classical chemical synapses. However, understanding of the properties and role of the former remains limited in the absence of sufficiently precise experimental tools. Here, we applied a novel strategy by attempting to convert a natural species difference in neuroendocrine dopamine (“TIDA”) neurons, which are strongly coupled in rats but completely lack electrical synapses in mice. By overexpressing the pore-forming protein, connexin-36, in mouse TIDA neurons, several measures of electrical coupling and synchronizing network effects were induced, albeit weakly. While network interactions were significantly altered by overexpression, overall circuit performance, reflected in target hormone levels, remained stable. These findings illustrate both the promise and current limitations of gain-of-function manipulations of electrical coupling.

## Introduction

Our ability to understand neural function relies in no small measure on elucidating the organizing principles and role of connectivity between nerve cells. For chemical synapses, such information has accumulated impressively with regard to both physiology and molecular composition over many decades. In contrast, electrical synapses have received notably less attention – a knowledge gap that does not reflect the importance of this mode of interneuronal communication (see Bennet MV, 1997). First described as a functional concept in crustaceans (Furshpan & Potter, 1957; Watanabe, 1958) and in structural terms in teleosts (M. V.L. Bennett et al., 1963), their existence was later demonstrated in the mammalian brain (Baker & Llinás, 1971). In the half-century since, electrical synapses, made up of gap junctions bridging the membranes of two neighbouring neurons, allowing for the passage of charge as well as signaling molecules and metabolites, have been shown to be a widespread phenomenon across phyla and neural circuits with great strategic importance for network function (see Alcamí & Pereda, 2019; Connors, 2017; Nagy et al., 2018; Söhl et al., 2005; Vaughn & Haas, 2022). The complex proposed effects of electrical coupling include *e.g.* synchrony (Michael V.L. Bennett & Zukin, 2004a; Long et al., 2004; Manor et al., 1997), anti-synchrony (Sherman & Rinzel, 1992; Terman et al., 2011), excitation (Bennett, 1997), inhibition (*e.g.* Gibson et al., 2005; Bennett, 1997; see also Vogelsy et al., 2013), and shunting (*e.g.* Lefler et al., 2014).

Progress in understanding electrical synapses has been hampered, however, by the paucity of tractable experimental tools. Pharmacological agents applied to block gap junction-mediated currents, *i.a.* 18-β-glycyrrhetinic acid (*e.g*. Guan et al., 2007), carbenoxolone (*e.g.* Beaumont & Maccaferri, 2011), octanol (*e.g*. Oku et al., 1999) and mefloquine (see Ghosh et al., 2021) are marred by off-target effects (see Juszczak & Swiergiel, 2009; Connors, 2012). Global genetic deletions of connexin 36 (Cx36), the main pore-forming component of neuronal electrical synapses (Condorelli et al., 1998), have aided in revealing *e.g.* the role of gap junctions in rhythmogenesis (Hormuzdi et al., 2001), but retain some residual interneuronal coupling (S. C. Lee et al., 2014), possibly by compensatory mechanisms (Bedner et al., 2012). While alternative strategies for interfering with electrical coupling by genetic introduction of point mutations in Cx36 have shown early promise (Placantonakis et al., 2006), the tool box for investigating electrical synapses in a physiological context remains lamentably limited.

The discovery, during the past decade, of an unexpected species difference in neuronal coupling offers a novel avenue for studying the role of gap junctions in the CNS, however. Thus, tuberoinfundibular dopamine (TIDA) neurons in the hypothalamus of the rat have been shown to be unusually strongly coupled by Cx36-containing electrical synapses, whereas such connections are completely absent in the mouse equivalent of this population, which lack Cx36 expression (Stagkourakis et al., 2018). The neuroendocrine TIDA neurons (Fuxe, 1965) play a central role in reproduction by serving as a “brake” on the pituitary production and release of the hormone, prolactin (see Grattan, 2015). The release of dopaminergic inhibition, most strikingly observed in mothers during the early postnatal period, underlies several parental functions, including lactation and care for offspring (see Qi-Lytle et al., 2023).

The network configuration of these neuroendocrine cells has powerful consequences at both the cellular and behavioural level. Thus, while the coupled rat TIDA neurons exhibit stereotyped, regular slow oscillations that are harmonized across the population (Lyons et al., 2010), mouse TIDA neurons exhibit significantly faster non-synchronous oscillations, in a range of frequencies (Stagkourakis et al., 2018). The frequency difference, in turn, has been shown to impact directly on parental behaviour, as male rats, as a consequence of their slow TIDA rhythms, have lower serum prolactin than mice and avoid contact with their pups, whereas mouse sires exhibit active pup care (Stagkourakis et al., 2020). Here, capitalizing on this species difference, we ask: to what extent can gap junctions be introduced in a normally uncoupled population, and what effect, if any, does that have for network operation and output? This question was addressed by introducing the expression of Cx36 in mouse TIDA neurons.

## Materials and Methods

### Animals

Male mice maintained on a C57BL/6J background were group-housed in a temperature- and humidity-controlled environment, with a 12/12-hour dark/light cycle (lights on at 6*am*) and *ad libitum* access to standard rodent chow and water. DAT-cre (Slc6a3^tm1(cre)Lrsn^; Ekstrand et al., 2007), tdTomato (strain 007914; Jackson Laboratories), and GCaMP6s (strain 028866, Jackson Laboratories) transgenic lines were maintained and cross-bred in-house. All experiments were approved by the local ethical committee and performed in accordance with the EU Directive 2010/63/EU, and the Swedish Board of Agriculture’s Regulations and General Advice on Laboratory Animals (SJVFS 2019:09, also known as L150).

### Stereotaxic surgery

Animals (4-7 weeks old) were kept under isoflurane vapor anaesthesia (5% for induction, 1.5%-2.5% maintenance). After anaesthesia induction, the animal was head-fixed in a stereotaxic frame (Stoelting or Neurostar). A small craniotomy was performed, and 300nl of AAV5-EF1a-DIO-Cx36-P2A-eYFP-WPRE-hGH (corresponding to the Cx36-OE group; 1,3 x 10^12^ viral copies/ml; Penn Vector core) or AAV5-EF1a-DIO EYFP (control group; 1,3 x 10^12^ viral copies/ml; Addgene) was injected through a glass capillary into the dmArc at the following coördinates from bregma: AP -1.80mm; L ±0.15mm; V -5.60mm, at a rate of 100nl/min. The plasmid vector pAAV-EF1a-DIO EYFP was purchased from Addgene (#27056) and was used to produce the custom-made AAV5-EF1a-DIO-Cx36-P2A-eYFP-WPRE-hGH (Penn Vector core). After an injection was completed, the capillary was left in place for 5mins, then retracted slowly. Injections were bi-or unilateral, as specified for each experiment in *Results*. Animals were allowed to recover, and electrophysiological recordings were performed at least three weeks after surgery.

### Acute slice preparation

Mice (8-15 weeks old) were deeply anaesthetised with isoflurane and decapitated. The brain was rapidly dissected and immersed in ice-cold, sucrose-based slicing solution (osmolarity 305-310mOsm), oxygenated with 95% O₂ / 5% CO₂, containing (in mM): sucrose (214), KCl (2), NaH_2_PO_4_ (1.2), NaHCO_3_ (26), glucose (10), MgCl_2_ (2), CaCl_2_ (2). Coronal slices containing the Arc (250μm thick) were obtained immediately using a vibratome (7000smz-2, Campden Instruments) and transferred into a bath of artificial cerebro-spinal fluid (aCSF; 305-310mOsm), containing (in mM): NaCl (127), KCl (2.0), NaH_2_PO_4_ (1.2), NaHCO_3_ (26), glucose (10), MgSO_4_ (1.3), CaCl_2_ (2.4), continuously bubbled with 95% O₂ / 5% CO₂. Slices were incubated at 34°C for 15mins for recovery, then kept at RT for at least 1h before experiments were performed. During electrophysiological recordings, the slices were continuously perfused with oxygenated aCSF at a rate of 2ml/min, in a temperature-controlled slice chamber equipped with an inline heater (Warner Instruments), and maintained at near-physiological temperature (34°C).

### Ex vivo electrophysiology

Electrophysiological recordings were obtained from TIDA neurons (identified as tdTomato-positive neurons in the dmArc), using the single- or paired whole-cell patch-clamp method. The slices were visualized using infrared differential interference contrast (DIC) imaging on an upright fluorescence microscope (Zeiss). Fluorescence filter cubes were used for visualization of tdTomato and eYFP fluorescence (Zeiss). In animals that had received AAV injections, only eYFP-positive TIDA neurons were selected for patch-clamp experiments. Patch pipettes (6-9MΩ) were filled with intracellular solution (280-290mOsm) containing (in mM): K-gluconate (140), KCl (10), HEPES (10), EGTA (1), MgCl_2_ (0.2), Na_2_ATP (2), Neurobiotin 0.2%, and pH adjusted to 7.3 with KOH. Recordings were performed in current-clamp mode, acquired in pClamp 10/11 software (Molecular Devices), using a MultiClamp 700B amplifier (Molecular Devices) and digitized at 20kHz using an Axon Digidata 1440B digitizer (Molecular Devices). Cells were recorded in gap-free mode for at least 10mins after the original stabilization period (*ca*. 5mins). Spontaneous activity was recorded first (no current injection) for 2-3mins, followed by injection of square negative step currents (-50pA to -10pA, 500ms long, > 1s intervals), and steady current holding values (-2pA to -30pA), to assess oscillation properties and electrical coupling (when applicable) at different cell membrane potential values. After the end of the recordings, each slice was returned to aCSF for at least 30mins before immersion-fixation in PFA 2%.

### Electrophysiology analysis

Data analysis was performed using ClampFit 11 software (Molecular Devices) and custom MATLAB scripts. Auto- and crosscorrelations were calculated using a custom MATLAB script from 60s of trace segments of spontaneous activity, after low-pass filtering (2Hz) to remove action potentials. Autocorrelation coefficient (ACC) is the highest detected peak in an autocorrelogram after t= 0s. The reported ACC_max_ corresponds to the membrane potential at which each cell exhibited the most stable oscillations by visual inspection (usually with maximum amplitude). The AC values of the five highest peaks of an autocorrelation function were arranged by their relative distance from zero, followed by data regression for each group (third-order polynomial curve fitting with least squares regression), as a proxy for the exponential decay of autocorrelation. Oscillation frequency was calculated as the reverse of the oscillation period identified from the autocorrelogram.

The crosscorrelation coefficient (CCC) denotes the peak of a crosscorrelogram with the greatest absolute magnitude, and lag denotes its temporal offset from zero. To identify whether a crosscorrelogram peak exceeds chance level, we applied block bootstrapping of 5s block size on both cell traces and computed the maximum crosscorrelation for 1000 iterations, creating a null distribution of CCC on the bootstrapped (shuffled) traces. If the detected CCC in the original traces exceeded the 5th-95th percentile of the null distribution in absolute value, it was considered significant. The coupling coefficient (CC) was calculated from negative step pulses during paired recordings. It is defined as the ratio of the response (*ΔVm*) of a non-manipulated cell (arbitrarily referred to as Cell 2) over the response (*ΔVm*) of the other, simultaneously recorded cell, which receives a current pulse (referred to as Cell 1), over the same period. Two time windows of 200ms were chosen, one just prior to a step pulse, and one at the end of a step pulse; the mean *Vm* of each cell before the step determines its baseline, whereas the mean *Vm* during the current injection determines its response. Input resistance was calculated from the same step pulse protocol, as the slope (*a*) of a linear regression model with the best fit (minimum Chi-square method) to the function *f(Vm)* = *aI*_input_, where *Vm* is the membrane potential of the cell, and *I*_input_ is the magnitude of the input current in pA.

### Ex vivo Ca^2+^ imaging and analysis

Ca^2+^ imaging experiments were performed in an electrophysiology setup, with the same acute slice preparation and maintenance during the experiment as for electrophysiology experiments (*vide supra*). The slices were imaged using a 20x water-immersion microscope objective (W Plan-Apochromat, 1.0 NA; Zeiss). Fluorescence videos of a cropped field-of-view containing the dmArc were captured at 20Hz (and downsampled by a factor of two) on a CMOS camera (Teledyne Technologies), acquired with the 4-D acquisition mode in Micro-Manager software (Edelstein et al., 2014). Typically, each slice was recorded for 10mins. The videos were processed using Inscopix Data Processing Software (Inscopix) to extract Ca^2+^ traces. The videos were down-sampled by a factor of two during analysis. Following pre-processing with a spatial bandpass filter (0.001– 0.5 pixel⁻¹) and motion-correction, the *ΔF/F* for each pixel was calculated as (*F – F_0_) / F_0_* (where the mean intensity of each corresponding pixel was used as *F_0_*). Lastly, regions of interest (ROIs) were manually drawn around all fluorescent cells in the dmArc, identified by visual inspection of maximal projections of the videos. Lastly, the ROI map was applied to the video, and Ca^2+^ traces were exported. Auto- and crosscorrelations were calculated from the entire traces using the same methodology and scripts as for electrophysiology recordings. Significant crosscorrelation was determined based on the bootstrapping method described in the electrophysiology analysis. Functional connectivity maps are graphs of all significantly crosscorrelated cell pairs, based on the X/Y location of their corresponding ROIs, and were created using a custom-made MATLAB script. Phase-locking values (PLVs) were calculated for all unique pairwise combinations. To determine significant phase-locking, a phase-randomization surrogate test was used as previously described (Lachaux et. al, 1999). Briefly, for each pairwise comparison, one trace was assigned random phase angles and the PLV with the experimental trace was calculated. A PLV null distribution was created from 1000 iterations of this process; pairs with experimental maximum PLV exceeding the 95th percentile of the null distribution were considered significantly phase-locked, and their average phase angle is reported. All network analyses were performed per hemisphere, and only unique pairwise comparisons are reported.

### Immunofluorescence

Following electrophysiology experiments, brain slices were immersion-fixed in 2% paraformaldehyde solution (PFA) for 16-20h, followed by a washing step in PBS for at least 1h with gentle agitation. Then, the slices were incubated in a blocking solution (10% standard donkey serum, 1% Bovine Serum Albumin, and 0.5% Tween 20 in PBS) for 30mins at RT, followed by incubation with primary antibodies (diluted in 1% standard donkey serum, 1% Bovine Serum Albumin and 0.5% Tween 20 in PBS) for 72h at 4°C with gentle agitation. At the end of the incubation period, the slices were washed three times for 5mins in PBS with 0.5% Tween 20, before the secondary, fluorophore-conjugated antibodies were applied for 2h at RT, in the dark, with gentle agitation. The same antibody diluent as for primary antibodies was used. The antibodies and dilution factors used in this study are as follows: anti-TH (AB_2737417; Encor; 1:1000), anti-GFP (immuno-reactive also against eYFP; AB_10000240; Aves Labs; 1:1000), and anti-Cx36 (AB_2533320; Invitrogen; 1:500) and secondary antibodies anti-rabbit-Alexa549, anti-Chicken-Alexa488, and anti-mouse-Cyanine5 (Invitrogen; 1:500). For visualization of neurons filled with Neurobiotin, Streptavidin-DyLight™405 (21831; Invitrogen; 1:500) was added to the first antibody incubation step, and slices were kept in the dark throughout the protocol. Following the secondary antibody incubation, the same washing steps were followed before the slices were transferred onto microscope glass slides and mounted using proLong™ Gold antifade mountant (Invitrogen). The samples were imaged on a confocal microscope (Airyscan 800; Zeiss or Stellaris 5; Leica) at least 16h later.

### Prolactin measurements (ELISA)

Trunk blood samples were collected during preparation of acute brain slices (*vide supra*). The samples were kept upright at RT for 30-60mins for the blood to clot, then centrifuged at 2000 RCF for 15mins at 4°C, on a pre-cooled centrifuge. The supernatant (serum) was collected, aliquoted, and stored at -80°C until further use. Prl concentrations were measured using a commercial sandwich ELISA kit (RAB0408; Merck-Millipore), on samples diluted 1:25 in assay buffer (as provided in the kit). Sample absorbance was measured, and a standard curve and median fluorescence intensity were obtained. Serum concentrations were inferred from a generated 4PL parameter logistic curve based on the given concentrations of the samples comprising the standard curve. Cx36-OE samples were from animals that had received bilateral injections with AAV5-EF1a-DIO-Cx36-P2A-eYFP-WPRE-hGH, whereas control animals received no injections. Duplicates of all samples were analysed with the mean value as the final reported concentration. Samples from animals used in electrophysiology and Ca^2+^ imaging experiments were included.

### Statistical Analysis

Statistical analysis was performed using GraphPad Prism 11 software. Normality was assessed for all datasets using D’Agostino-Pearson and Shapiro-Wilk tests, and appropriate parametric or non-parametric tests were performed, typically unpaired t-test with Welch’s correction (referred to as Welch’s test) or Mann-Whitney test. Contingency tables were computed using two-sided Fisher’s exact test. Polynomial regression curves were compared using Akaike’s Information Criterion (AIC) to estimate differences in data fitting. The plots represent the median with a horizontal line and the 1st-3rd quartile range with dotted lines, unless otherwise described. Whiskers represent the minimum and maximum values, unless otherwise described. All statistical tests performed are described in the figure legends. Representation of significance is marked for all plots as follows: non-significant P≥ 0.05 (ns), P<0.05 (*), P<0.01 (**), P<0.001 (***), P<0.0001 (****).

## Results

### Transduction of Cx36 in mouse TIDA neurons leads to sparse electrical coupling

First, we set out to establish whether transferring the Cx36-encoding *Gjd2* gene in TIDA neurons leads to detectable Cx36 signal in the cell membrane. Male DAT:cre-tdTomato mice received stereotaxic injections in the dmArc of the Cx36-overexpression construct AAV5-EF1a-DIO-Cx36-P2A-eYFP-WPRE-hGH (“Cx36-OE mice”). Transfected neurons in these animals exhibited widespread Cx36-immunoreactivity (IR). In contrast, control animals injected with the same viral construct absent the Cx36 gene, showed high levels of the reporter eYFP-IR, but no Cx36 signal (Fig. 1A-C). Presence of eYFP was restricted to dopaminergic cells (TH-positive) in both groups (Fig. 1A-C), verifying Cre-dependent expression of the AAV. These data suggest that introduction of the transgene into TIDA neurons generates the molecular substrate for GJs in these cells.

**Figure 1.**
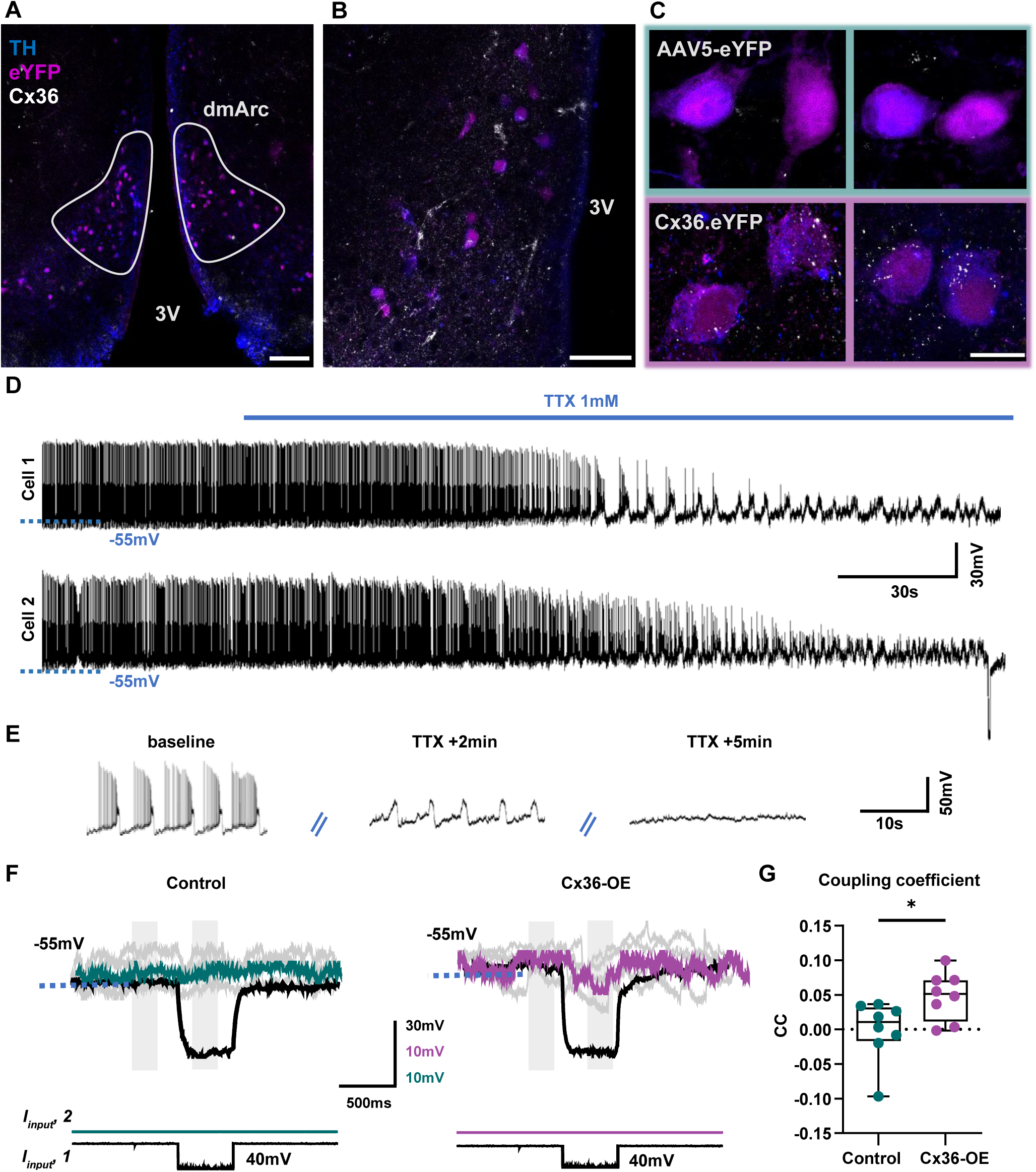
Transduction of connexin-36 (Cx36) leads to weak electrical coupling in mouse TIDA neurons. ***A***, Confocal micrograph from a section containing the dorsomedial arcuate nucleus (dmArc; outlined in white) processed with immunofluorescence for tyrosine hydroxylase (TH; blue), enhanced yellow fluorescent protein (eYFP; magenta), and Cx36 (white), from a mouse virally transduced with the Cx36 construct. Note clustering of TIDA neurons in the dmArc adjacent to the third ventricle (3V). Scale bar, 100μm. ***B,*** Cx36 immunoreactivity on transfected TIDA neurons of a Cx36-ΟΕ animal, raw signal. Scale bar, 20μm. ***C***, High-resolution micrographs of TIDA neurons from an animal injected with AAV5-eYFP virus (control group, top, teal box) or Cx36.eYFP virus (Cx36-OE group, bottom, purple box). Note presence of Cx36 immunoreactivity (white puncta) in the latter but not the former. Scale bar: 10 μm. ***D,*** Example trace of paired whole-cell patch-clamp recording of TIDA neurons and the time course of the 1 mM tetrodotoxin (TTX) effect. Note gradual dampening of phasic firing ***E***, Traces from cells shown in *D,* visualized at greater temporal resolution. Note the rhythmic spontaneous baseline oscillations which persist after action potential blockade by TTX (2mins of application) and eventually disappear after longer (5mins) TTX application. ***F***, Example traces of voltage responses to negative square current pulses, during paired recordings in control (left) and Cx36-OE (right) neurons, in the presence of TTX. Recorded individual responses (grey traces) and response averaged over five consecutive sweeps in black (Cell 1; pulse-injected cell) or teal/purple (Cell 2; no direct current injection). Teal indicates control TIDA neurons, purple indicates Cx36-OE neurons. Grey boxes highlight the 200ms periods before and during the end of a step response, respectively, from which the coupling coefficient is calculated. ***G,*** Coupling coefficient measured under application of TTX is higher in Cx36-OE neuron pairs recorded by patch-clamp (n_control_= 4 pairs from 3 animals, n_Cx36-OE_ 4 pairs from 4 animals; Mann-Whitney test).

Next, paired whole-cell patch-clamp recordings were performed on acute brain slices to assess electrical coupling. To directly evaluate coupling, we examined subthreshold voltage transfer during square pulse current injections (see *Methods*). However, in these spontaneously oscillating neurons, baseline membrane potential (V_m_) fluctuates significantly with high frequency, which may confound such measurements. Application of the voltage-dependent Na^+^ channel antagonist, tetrodotoxin (TTX, 1 μM), has been shown to abolish or dampen mouse TIDA oscillations (Stagkourakis et al., 2018). Thus, TTX was applied for a minimum of 5mins to stabilize V_m_ as much as possible, abolishing oscillatory activity (Fig. 1D-E). In control pairs, no electrical coupling was observed (with CC of -0,0008 ± SD 0.043; Fig. 1F-G). In contrast, CC was predominantly positive (0.048 ± SD 0.035) and of smaller variance than in controls, in the Cx36-OE group (P =0.0160, U = 9.50, Mann-Whitney; Fig. 1F-G;). These data suggest that a degree of electrical coupling has been established by the OE paradigm, though substantially lower than that observed in the naturally Cx36-expressing rat TIDA system (Stagkourakis et al., 2018).

Motivated by these results, we wanted to examine coupling under naturalistic conditions as well. While TTX aids in achieving a more stable V_m_, this pharmacological manipulation also removes part of the TIDA membrane property repertoire as the persistent Na^+^ current appears to contribute to rhythm generation in rodent TIDA neurons (Lyons et al., 2010). Thus, step protocols were repeated, but now in the absence of TTX. Typically, rhythmic activity across paired TIDA neurons appeared asynchronous in both control and Cx36-OE mice (Fig 2A). In control pairs, where, on average, coupling was absent (-0.021 ± SD 0.046; Fig. 2B, C), negative CC values were often observed, reflecting a V_m_ *increase* in the non-manipulated cell and underscoring the interference of the fluctuating baseline in spontaneously active TIDA neurons. While directly observable electrical coupling was rare in the Cx36-OE group, such pairs exhibited, on average, significantly higher CC than controls (P= 0.0079, U= 90.50, Mann-Whitney; Fig. 2B-C) with a noticeably smaller representation of negative values. Interestingly, the maximum crosscorrelation coefficient (CCC_max_) of sub-threshold oscillations was significantly higher among Cx36-OE pairs (P =0.0037, R^2^ =0.418, Welch’s test; Fig. 2D), suggesting that rhythms are coördinated by some mechanism in this group. Furthermore, in recordings where one neuron had been filled with Neurobiotin, a few occasions of dye-coupling were observed (Fig. 2E). Together, these data provide further support the existence of sparse induced electrical coupling across the Cx36-OE TIDA population.

**Figure 2.**
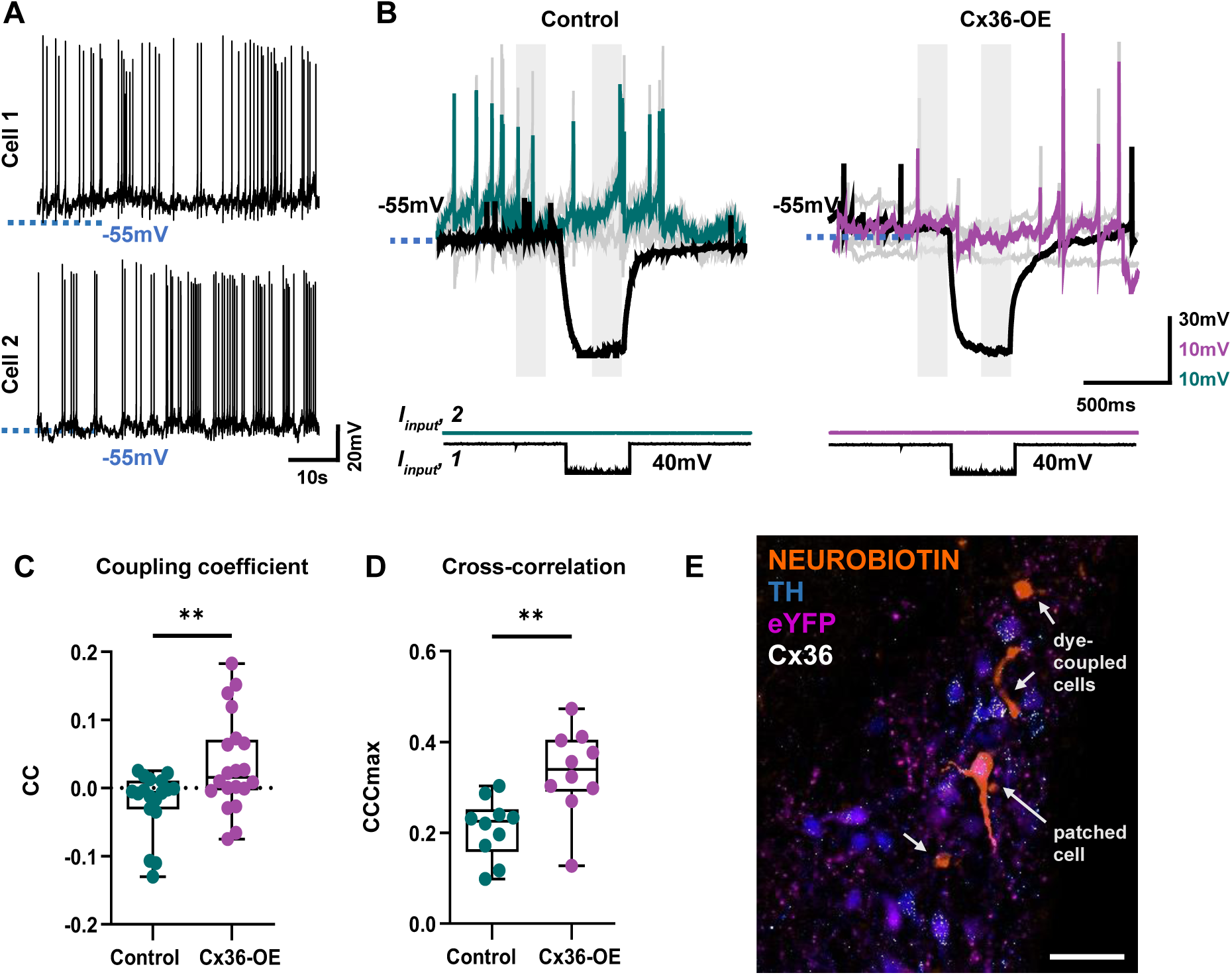
Electrical coupling during spontaneous TIDA oscillations. ***A***, Representative example of a paired recording showing spontaneous, asynchronous oscillations observed in TIDA neurons. This particular example was acquired from a Cx36-overexpressing (Cx36-OE) pair. ***B***, Example traces of responses to pulse currents during spontaneous paired recordings in control (left) and Cx36-OE (right) neurons, organized as in Fig 1F. ***C***, ***D***, Coupling coefficient (***C***) and crosscorrelation (***D***), recorded from control (teal) and Cx36-OE (purple) neuron pairs (n_control_= 9(***C***)/10(***D***) pairs from 7 animals, n_Cx36-OE_= 10 pairs from 9 animals; (***C***) Mann-Whitney test; (***D***) Welch’s test). Note significantly higher averages for both parameters in the latter group. ***E***, Confocal micrograph of a slice containing the dorsomedial arcuate nucleus, used for whole-cell patch-clamp electrophysiology using a Neurobiotin (NB)-containing recording micropipette, *post-hoc* processed with immunofluorescence for NB (orange), tyrosine hydroxylase (TH; blue), enhanced yellow fluorescent protein (eYFP; magenta), and connexin-36 (Cx36; white). Note dye coupling; patch-clamped TIDA neuron indicated by the long arrow; short arrows point to three instances of non-recorded cells exhibiting NB fluorescence, reflecting likely dye coupling. Scale bar: 50μm.

### Similar oscillation properties with enhanced rhythmicity in the Cx36-OE TIDA population

Next, we addressed whether the imposed expression of Cx36 affected TIDA oscillation dynamics. Both control and Cx36-OE TIDA neurons exhibited heterogeneous activity patterns (activity traces in Fig. 3A-B), as previously reported. Oscillation frequency (P=0.1219, U= 524.5, Mann-Whitney; Fig. 3C) and passive membrane properties, such as input resistance (P=0.2445, R^2^ =0.0362; Welch’s test; Fig. 3D), were similarly distributed between groups. However, Cx36-OE neurons exhibited a significantly increased V_m_ maximum autocorrelation coefficient (ACC_max_; P =0.0158, U = 447.5, Mann-Whitney; Fig. 3E). Notably, Cx36-OE TIDA neurons demonstrated several autocorrelation peaks near the ACC_max_ value, and their autocorrelograms were characterized by slower oscillation dampening (P< 0.0001, AIC difference= 13.60, Akaike’s Information Criterion comparison of fits; Fig. 3F), reflecting more rhythmic and stable oscillations, presumably maintained through electrical coupling.

**Figure 3.**
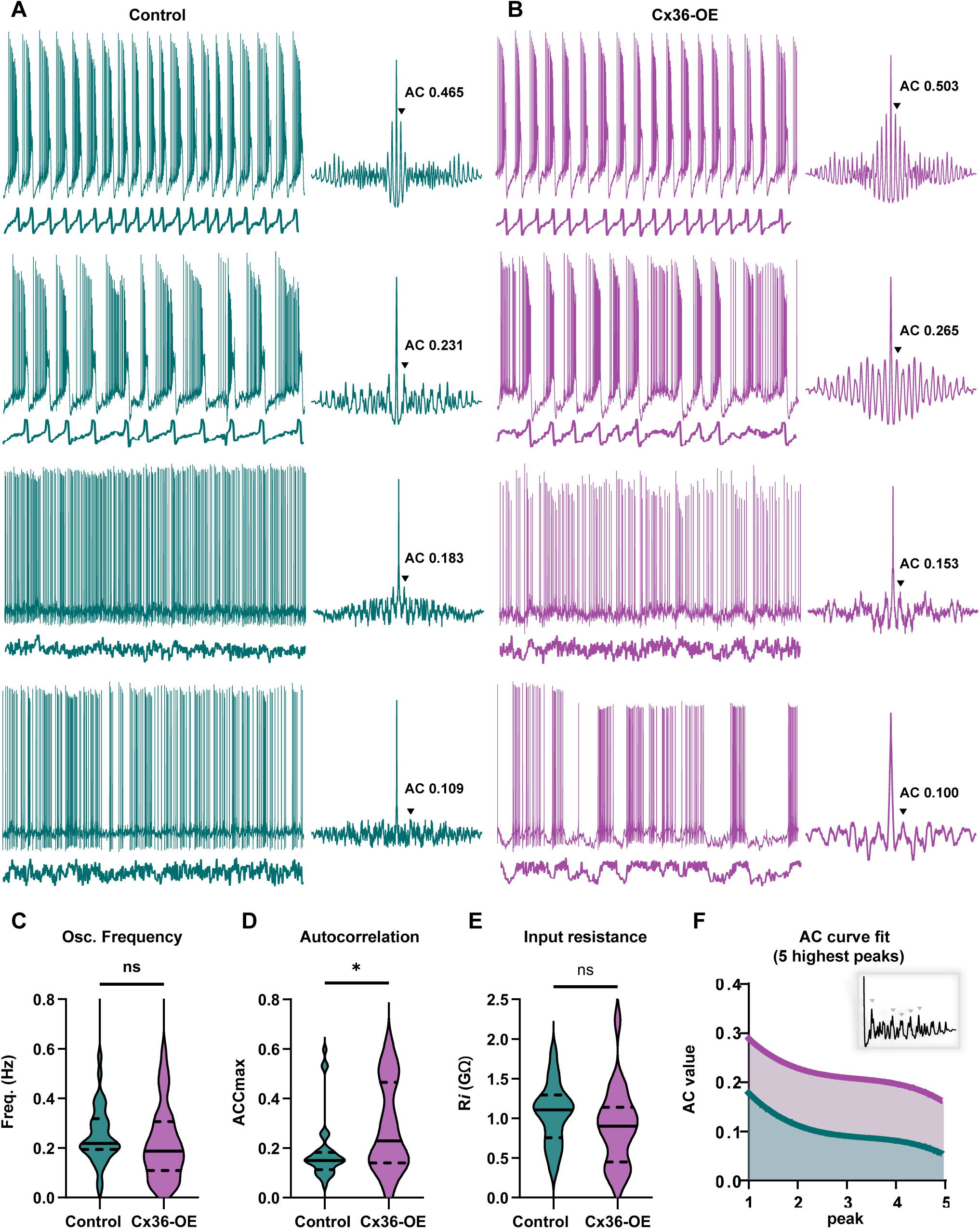
TIDA oscillation properties under control and connexin-36 overexpression (Cx36-OE) conditions. ***A, B,*** Example traces and autocorrelation curves of different types of oscillatory activity observed in control (***A***; teal) and Cx36-OE (***B***; purple) TIDA neurons. Top traces show unfiltered recordings; bottom traces show same recording low-pass filtered at 2Hz to reveal the underlying membrane potential pattern in the absence of action potentials. Black arrowheads indicate the maximal autocorrelation coefficient (ACC_max_). ***C***, ***D***, Oscillation frequency (***C***) and input resistance (***D***), recorded from control and Cx36-OE neurons (***C***: n_control_= 35 cells from 11 animals, n_Cx36-OE_ = 38 cells from 19 animals; Mann-Whitney test. ***D***: n_control_= 19 cells from 10 animals, n_Cx36-OE_ = 21 cells from 17 animals; Welch’s test); neither parameter is significantly different between the groups. ***E***, ACC_max_ calculated from control and Cx36-OE TIDA neurons, during 60s of spontaneous activity (n_control_= 35 cells from 11 animals, n_Cx36-OE_ = 38 cells from 19 animals); note higher ACC_max_ in Cx36-OE group. ***F***, Cubic polynomial curve fitting of the five highest peaks of the autocorrelation function of control cells (teal) and Cx36-OE cells (purple). Note the slower decay of the fitted curve in Cx36-OE cells (P<0.0001, Akaike’s Information Criterion comparison of fits). The insert shows an example autocorrelogram with the five highest peaks marked.

### Increased TIDA functional connectivity and in-phase oscillations within the network

The higher prevalence of stable oscillations, and the stronger crosscorrelation observed in paired Cx36-OE recordings may reflect broader differences in coördination of TIDA network dynamics. To capture the activity relationships on a network-wide level, widefield Ca^2+^ imaging was performed on acute brain slices of DAT:cre-GCaMP6s male mice. Ca^2+^ activity traces and functional connectivity maps were computed after applying block-bootstrapping (see *Materials and Methods)* to identify significantly correlated pairs (Fig. 4A-B). Notably, networks of the Cx36-OE model showed higher functional connectivity, with more abundant significantly correlated pairs (P=0.0011, Fisher’s exact test, all lags considered; Fig. 4C). When examining pairs with shorter temporal offsets (*e.g*., those with lags < 2.5 s– *i.e*., half the average oscillation period), the CCC tends to be higher in Cx36-OE networks. Interestingly, this difference is significant for pairs of neurons closely in-phase (positive CCC), whereas pairs close to anti-phase (negative CCC) did not show this discrepancy (Fig. 4D-F; in-phase: U = 41.50, P =0.0130, anti-phase: U = 82.50, P =0.3568; Mann-Whitney test).

**Figure 4.**
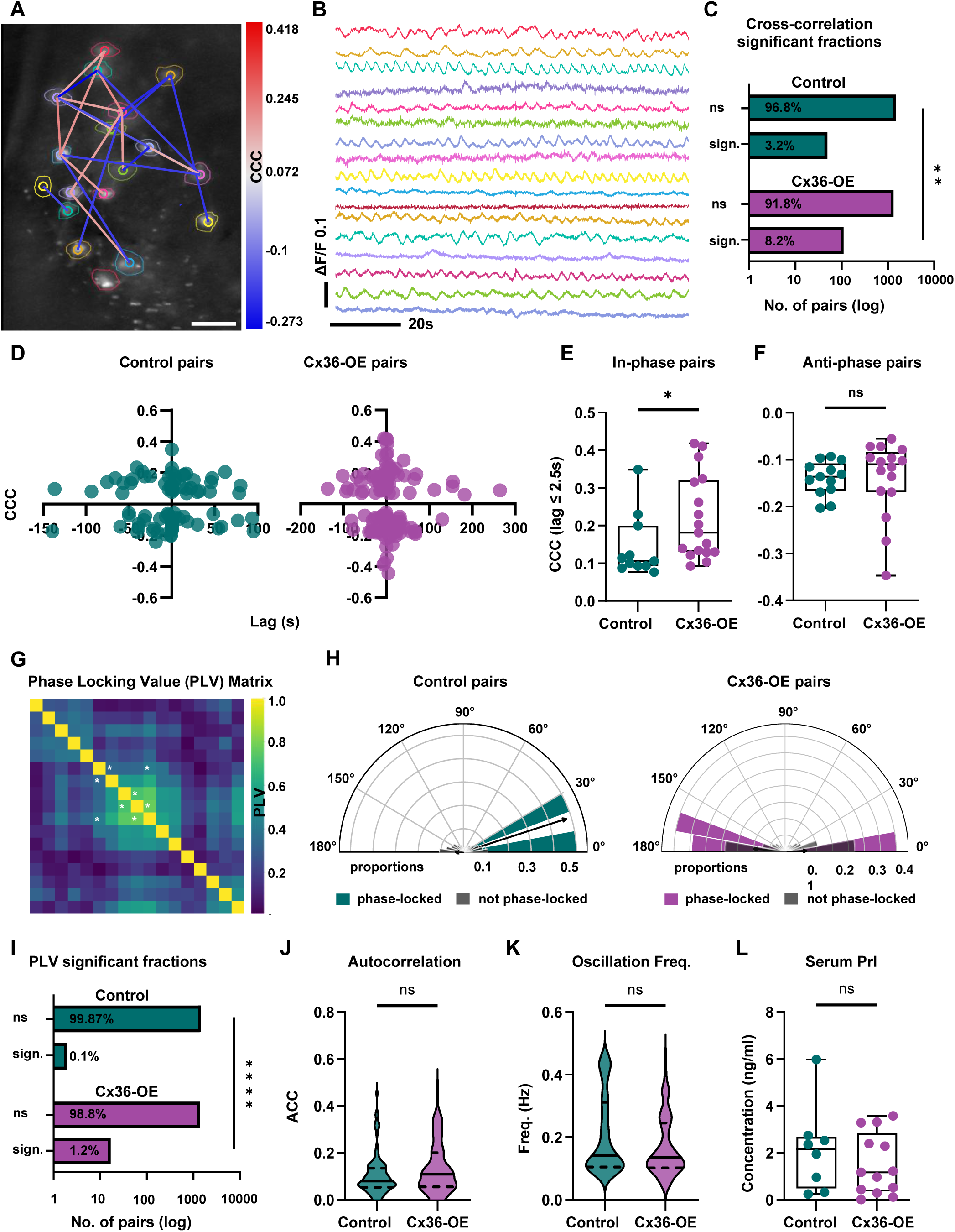
Connexin-36 overexpression (Cx36-OE) results in changes in network dynamics but not prolactin levels. ***A***, Example micrograph of a dmArc slice from DAT-cre:GCaMP6s male mouse. TIDA neurons are color-coded as manually identified regions of interest with their computed functional connectivity map, as deduced from Ca^2+^ imaging, superimposed. Only significant connections are represented. Heatmap to the right represents the crosscorrelation coefficient (CCC) for each pair. Scale bar: 20 μm. ***B***, Extracted Ca^2+^ traces from the same regions of interest as in (***A***), showing a range of oscillatory dynamics. ***C,*** Contingency plots for the log-fractions of significantly (sign.) and non-significantly (ns) crosscorrelated pairs (P=0.0011, Fisher’s exact test). The corresponding percentage fractions are indicated on each bar. Note the significantly higher fraction of functionally connected pairs among Cx36-OE networks. ***D***, Crosscorrelation coefficients of all significantly correlated cell pairs, and their respective lag values, across all control (left) and Cx36-OE (right) networks. ***E***, ***F***, Crosscorrelation coefficients of pairs with close temporal offsets (significantly correlated and with lag ≤ 2.5s). ***E***, Positive CCC values indicate in-phase pairs (n_control_= 11 cells from 7 animals, n_Cx36-OE_ = 17 cells from 6 animals; Mann-Whitney test). Note higher CCC values in Cx36-OE pairs. ***F***, Negative CCC values indicate anti-phase pairs (n_control_= 13 cells from 7 animals, n_Cx36-OE_ = 16 cells from 6 animals; Mann-Whitney test). ***G***, Example Phase Locking Value (PLV) matrix heatmap with four directional pairs exhibiting significant PLV (white asterisks). ***H***, Histograms of the calculated average phase angles for all pairwise comparisons in control (left) and Cx36-OE (right) networks. The respective proportions of non-significantly (grey) and significantly phase-locked pairs in control (teal) and Cx36-OE (purple) are indicated below the angle plots. Black vector arrows mark the population average. ***I***, Contingency plot for the log-fractions of significantly (sign.) and non-significantly (ns) phase-locked pairs (P=0.0003, Fisher’s exact test). The corresponding percentage fractions are indicated on each bar. Note the significantly higher fraction of phase-locked pairs among Cx36-OE networks. ***J, K***, Autocorrelation coefficients (ACCs) and oscillation frequencies calculated from the entire Ca^2+^ traces across all recorded cells (n_control_= 98 cells from 7 animals, n_Cx36-OE_ = 81 cells from 6 animals; Mann-Whitney tests). Note that both properties are similar between the groups. ***L***, Prolactin concentration in serum, determined by ELISA. Note that concentrations are not significantly different between the groups (n_control_= 8 mice, n_Cx36-OE_= 13 mice; Welch’s test).

Furthermore, we examined the consistency of phase relationships by calculating pairwise phase-locking values (PLVs) and phase angles (Fig. 4G-H). Although a small fraction of the control mouse TIDA population is phase-locked (0.12%), a significantly larger fraction (1,70%) is observed in the Cx36-OE group (P= 0.0003, Fisher’s exact test; Fig. 4H-I). Markedly, phase-locked pairs oscillate mostly in-phase or anti-phase (*i.e*., with phase angles close to 0° or 180°, respectively), but no intermediate phase relationships are observed (Fig. 4H). Lastly, ACCs and oscillation frequencies computed from the entire traces (*ca*. 10 mins) were distributed similarly between the groups (ACC: P = 0.0752, U= 3181; oscillation frequency: P =0.1015, U= 3004; Mann-Whitney; Fig. 4J-K), indicating that oscillation frequency, unlike rhythmicity, is closely maintained at longer time scales. These data suggest stronger functional connectivity with a higher degree of oscillation synchrony and phase-locking when Cx36 expression is imposed, both hallmarks of the presence of electrical coupling within a network (Steriade, 1997; Bennett and Zukin, 2004; Terman et.al., 2011).

### Network output is preserved

TIDA oscillation frequency has been causally implicated in DA release and, consequently, circulating Prl levels (Stagkourakis et al., 2020). Given our observations of altered coupling and network dynamics in Cx36-OE male mice, we determined serum Prl concentration through ELISA. Peripheral Prl levels were not significantly different between controls and bilaterally transduced Cx36-OE mice (R^2^ =0.06, P =0.4156, Welch’s test; Fig. 4L). Thus, the observed changes in TIDA cellular and network properties do not appear to be sufficient to perturb set points within the male mouse lactotrophic axis.

## Discussion

Electrical synapses are now established as broadly distributed in the nervous system across the animal kingdom, and important for neural circuit operations (Michael V.L. Bennett & Zukin, 2004b; Harris, 2018). However, their exact role, especially within larger networks, remains poorly understood due to limitations in specificity and target precision of available experimental tools. Here, we apply a novel strategy by introducing Cx36 expression in a natively uncoupled cell population for which a close coupled correlate exists in a related species; *i.e.* by virally mediated transduction into mouse TIDA neurons, which, unlike in rats, do not establish electrical synapses (Stagkourakis et al., 2018).

Our genetic manipulation yielded robust and specific expression of Cx36 immunoreactivity in TIDA neurons. The presence of protein signal alone does not guarantee the functional assembly and insertion of gap junctions. However, the hallmark of electrical synapses, *i.e.* the subthreshold voltage transfer between neighbouring neurons, which is normally completely absent from the mouse TIDA system (Stagkourakis et al., 2018), was detected in a large proportion of recorded Cx36-OE neuron pairs. Electrical CC was typically on the order of 0.07. This is notably lower than the rat TIDA system, where the average recorded CC is 0.18 (Stagkourakis et al., 2018). Thus, a full conversion to the gap junction-linked rat TIDA system was not achieved. However, in a brain-wide perspective, it should be noted that the latter number is among the highest reported in any CNS system (see Curti et al., 2012). Furthermore, in the inferior olivary nucleus, perhaps the best studied example of electrical synapses in the mammalian brain, typical CC <0.05 is reported (Devor & Yarom, 2002) Such numbers have been reported from *i.a.* interneuron networks in the hippocampus (*e.g.* Zsiros & Maccaferri, 2005), and in the neocortex (*e.g.* Galarreta & Hestrin, 1999); and even weaker in adulthood;(Galarreta & Hestrin, 2002), the thalamic reticular nucleus (*e.g.* Landisman et al., 2002), cerebellar Golgi cells (*e.g.* Brunel et al., 2009) and the suprachiasmatic nucleus (*e.g.* Long et al., 2004). Thus, the electrical coupling that was achieved by artificially driving Cx36 expression in a natively uncoupled system (present study) was on the order of what is physiologically observed in many brain regions (see Haas, 2015).

At the cellular and network levels, Cx36-OE cells exhibited higher measures of cross-correlation and oscillations that were both more rhythmic and more stable than in TIDA neurons from control mice. Functional connectivity and synchrony were also higher in Cx36-OE mice when studied at a population level. Notably, in these animals, oscillatory phase relationships clustered towards the extremes of in-phase (0°) or antiphase (180°), but not in between (Fig. 4H). This phenomenon has been observed in computational models of weakly electrically coupled oscillating neurons (E. Lee & Terman, 2013; Sharp et al., 1992; Sherman & Rinzel, 1992; Terman et al., 2011). Together, these findings further support the conclusion that Cx36 expression resulted in functional gap junctions between TIDA neurons.

Yet, while Cx36-OE led to significant changes at the network level, this manipulation did not alter the dopamine-mediated inhibition of pituitary hormone release. This was evidenced by unaltered circulating prolactin levels in Cx36-OE mice, which did not match the low concentrations observed in male rats (Stagkourakis et al., 2020). Indeed, average oscillation frequency – a parameter that has been causally implicated as determining dopamine output (Stagkourakis et al., 2020) – did not change with the genetic introduction of connexin. Furthermore, the near-complete oscillation synchrony observed in the rat TIDA network (Lyons et al., 2010; Stagkourakis et al., 2018) was not observed in Cx36-OE networks. Thus, while the present data provide evidence for successful exogenous introduction of gap junctions in a natively uncoupled system, they also highlight the limitations of this manipulation in altering network configuration and output.

What could be the constraining factors that impede achieving a more impactful, network-wide coupling? Firstly, it should be kept in mind that while connexins are the indispensable pore-forming component of gap junctions, electrical synapses also include, and depend on, other proteins. Chief among these is the scaffolding protein, Zona Occludens-1, which interacts with connexins in gap junctions (Giepmans & Moolenaar, 1998), and, critically, with Cx36 in neurons (Lasseigne et al., 2021; Li et al., 2004), but there is also a growing list of other proteins implicated in stabilizing the intercellular connection (see Miller & Pereda, 2017). To what extent such components are available in mouse TIDA neurons, and their requirement for gap junction assembly, membrane insertion, trafficking, and recycling remains to be established. Secondly, cell membranes need to be closely apposed for the intercellular channels of electrical synapses to form. Typical distances reported in ultrastructural studies of gap junctions are in the range of 2-4 nm (Brightman & Reese, 1969; Goodenough & Revel, 1970; Makowski et al., 1977). Although appositions of neighbouring cell somata, and sparse physical dendritic connections or crossings, are seen on occasion among murine TIDA neurons (our own unpublished observations), it is not known at present what the membrane distances are in the mouse (or rat) TIDA systems, or on which cell compartments (*e.g.* somata, dendrites) electrical synapses localize on these neurons. Finally, unlike in the rat, where gap junctions have presumably been present from (or before) birth in the TIDA system, Cx36 was introduced in mouse TIDA neurons after weaning, in adolescence, in our study. It is possible that electrical synapses, which in many systems are at their most prevalent during development with a subsequent decrease (Belousov, 2011; M. V.L. Bennett et al., 1981), need to be established early in life to be fully functional.

The current results should be interpreted with both caution and restrained optimism. The introduction of Cx36 *in vivo* successfully achieved strong expression of the protein and coupling strength on par with many other brain regions. Yet, to the extent that the rat can serve as a “ground truth” for an electrically coupled TIDA system, the network configuration and output of that species were not observed, reflecting the current limitations of this approach. Notably, however, a recent study (Ransey et al., 2026) elegantly introduced constructs of engineered connexins in murine local cortical interneuron networks and a long-range corticothalamic circuit to successfully enhance oscillatory coupling and influence circuit-specific behaviour. Other studies in the literature on imposed coupling in neural cells through Cx overexpression have been restricted to cell cultures (e.g. Srinivas et al., 1999; Bargiello et al., 2019). Future work, exploring underlying mechanisms and further refinements, will show the potential for exploiting gap junction manipulation as a means for circuit editing in the nervous system.

## Author contributions

A.M. and C.B. designed research; A.M. performed research; A.M. and A.L. analyzed data; A.M., A.L. and C.B. wrote the manuscript; C.B. acquired funding

## Acknowledgements

The work described here was made possible through the generous support of a European Research Council Advanced Grant (TOGETHER; Grant Agreement No. 101021496) and the Swedish Research Council Distinguished Professor Program (2021-00671_VR) to CB, and internal funds from Stockholm University.

We thank Stockholm University’s Experimental Core Facility (ECF), as well as the Imaging Facility at Stockholm University (IFSU) for assistance with confocal microscopy. Furthermore, we thank Dr. Paul Williams and Dr. Maria Jimena Ferraris for their assistance in animal colony management and technical assistance.

## Competing interes

The authors disclose no competing interests.

## References

Alcamí, P., & Pereda, A. E. (2019). Beyond plasticity: the dynamic impact of electrical synapses on neural circuits. Nature Reviews Neuroscience, 20(5), 253–271. 10.1038/s41583-019-0133-5

Baker, R., & Llinás, R. (1971). Electrotonic coupling between neurones in the rat mesencephalic nucleus. The Journal of Physiology, 212(1), 45–63. 10.1113/JPHYSIOL.1971.SP009309

Bargiello, T. A., Oh, S., Tang, Q., Bargiello, N. K., Terry, L., Kwon, T., & States, U. (2019). HHS Public Access. 1860(1), 22–39. 10.1016/j.bbamem.2017.04.028.Gating

Beaumont, M., & Maccaferri, G. (2011). Is connexin36 critical for GABAergic hypersynchronization in the hippocampus? The Journal of Physiology, 589(Pt 7), 1663. 10.1113/JPHYSIOL.2010.201491

Bedner, P., Steinhäuser, C., & Theis, M. (2012). Functional redundancy and compensation among members of gap junction protein families? Biochimica et Biophysica Acta-Biomembranes, 1818(8), 1971–1984. 10.1016/j.bbamem.2011.10.016

Belousov, A. B. (2011). The regulation and role of neuronal gap junctions during development. Communicative & Integrative Biology, 4(5), 579–581. 10.4161/CIB.16380

Bennett, M. V.L., Aljure, E., Nakajima, Y., & Pappas, G. D. (1963). Electrotonic junctions between teleost spinal neurons: Electrophysiology and ultrastructure. Science, 141(3577), 262–264. 10.1126/SCIENCE.141.3577.262

Bennett, M. V.L., Spray, D. C., & Harris, A. L. (1981). Electrical Coupling in Development. Integrative and Comparative Biology, 21(2), 413–427. 10.1093/ICB/21.2.413

Bennett, Michael V.L., & Zukin, R. S. (2004a). Electrical Coupling and Neuronal Synchronization in the Mammalian Brain. Neuron, 41(4), 495–511. 10.1016/S0896-6273(04)00043-1

Bennett, Michael V.L., & Zukin, R. S. (2004b). Electrical Coupling and Neuronal Synchronization in the Mammalian Brain. Neuron, 41(4), 495–511. 10.1016/S0896-6273(04)00043-1

Brightman, M. W., & Reese, T. S. (1969). Junctions between intimately apposed cell membranes in the vertebrate brain. The Journal of Cell Biology, 40(3), 648–677. 10.1083/jcb.40.3.648

Brunel, N., Hakim, V., Schwartz, E., Chat, M., Le, M., Dugue, G. P., & Courtemanche, R. (2009). Article Electrical Coupling Mediates Tunable Low-Frequency Oscillations and Resonance in the Cerebellar Golgi Cell Network. 126–139. 10.1016/j.neuron.2008.11.028

Condorelli, D. F., Parenti, R., Spinella, F., Salinaro, A. T., Belluardo, N., Cardile, V., & Cicirata, F. (1998). Cloning of a new gap junction gene (Cx36) highly expressed in mammalian brain neurons. European Journal of Neuroscience, 10(3), 1202–1208. 10.1046/J.1460-9568.1998.00163.X;REQUESTEDJOURNAL:JOURNAL:14609568;WGROUP:STRING:PUBLICATION

Connors, B. W. (2012). Tales of a Dirty Drug: Carbenoxolone, Gap Junctions, and Seizures: Carbenoxolone and Seizures. Epilepsy Currents, 12(2), 66–68. 10.5698/1535-7511-12.2.66

Connors, B. W. (2017). Synchrony and so much more: Diverse roles for electrical synapses in neural circuits. Developmental Neurobiology, 77(5), 610–624. 10.1002/dneu.22493

Curti, S., Hoge, G., Nagy, J. I., & Pereda, A. E. (2012). Synergy between Electrical Coupling and Membrane Properties Promotes Strong Synchronization of Neurons of the Mesencephalic Trigeminal Nucleus. The Journal of Neuroscience, 32(13), 4341. 10.1523/JNEUROSCI.6216-11.2012

Devor, A., & Yarom, Y. (2002). Electrotonic coupling in the inferior olivary nucleus revealed by simultaneous double patch recordings. Journal of Neurophysiology, 87(6), 3048–3058. 10.1152/jn.2002.87.6.3048

Edelstein, A. D., Tsuchida, M. A., Amodaj, N., Pinkard, H., Vale, R. D., & Stuurman, N. (2014). Advanced methods of microscope control using µ Manager software. 1(2), 1–10. 10.14440/jbm.2014.36

Ekstrand, M. I., Terzioglu, M., Galter, D., Zhu, S., Hofstetter, C., Lindqvist, E., Thams, S., Bergstrand, A., Hansson, F. S., Trifunovic, A., Hoffer, B., Cullheim, S., Mohammed, A. H., Olson, L., & Larsson, N. G. (2007). Progressive parkinsonism in mice with respiratory-chain-deficient dopamine neurons. Proceedings of the National Academy of Sciences of the United States of America, 104(4), 1325–1330. 10.1073/PNAS.0605208103

Furshpan, E. J., & Potter, D. D. (1957). Mechanism of Nerve-Impulse Transmission at a Crayfish Synapse. Nature 1957 180:4581, 180(4581), 342–343. 10.1038/180342a0

Fuxe, K. (1965). Evidence for the existence of monoamine neurons in the central nervous system. Zeitschrift Für Zellforschung Und Mikroskopische Anatomie 1965 65:4, 65(4), 573–596. 10.1007/BF00337069

Galarreta, M., & Hestrin, S. (1999). A network of fast-spiking cells in the neocortex connected by electrical synapses. Nature, 402(6757), 72–75. 10.1038/47029

Galarreta, M., & Hestrin, S. (2002). Electrical and chemical synapses among parvalbumin fast-spiking GABAergic interneurons in adult mouse neocortex. Proceedings of the National Academy of Sciences of the United States of America, 99(19), 12438–12443. 10.1073/PNAS.192159599

Ghosh, D. K., Kumar, A., & Ranjan, A. (2021). Cellular targets of mefloquine. Toxicology, 464, 152995. 10.1016/J.TOX.2021.152995

Gibson, J. R., Beierlein, M., & Connors, B. W. (2005). Functional properties of electrical synapses between inhibitory interneurons of neocortical layer 4. Journal of Neurophysiology, 93(1), 467–480. 10.1152/JN.00520.2004

Giepmans, B. N. G., & Moolenaar, W. H. (1998). The gap junction protein connexin43 interacts with the second PDZ domain of the zona occludens-1 protein. Current Biology, 8(16), 931–934. 10.1016/S0960-9822(07)00375-2

Grattan, D. R. (2015). 60 YEARS OF NEUROENDOCRINOLOGY: The hypothalamo-prolactin axis. The Journal of Endocrinology, 226(2), T101–T122. 10.1530/JOE-15-0213

Guan, B. C., Si, J. Q., & Jiang, Z. G. (2007). Blockade of gap junction coupling by glycyrrhetinic acids in guinea pig cochlear artery: A whole-cell voltage- and current-clamp study. British Journal of Pharmacology, 151(7), 1049. 10.1038/SJ.BJP.0707244

Haas, J. S. (2015). A new measure for the strength of electrical synapses. Frontiers in Cellular Neuroscience, 9(September), 378. 10.3389/FNCEL.2015.00378

Harris, A. L. (2018). Electrical coupling and its channels. Journal of General Physiology, 150(12), 1606–1639. 10.1085/jgp.201812203

Hormuzdi, S. G., Pais, I., LeBeau, F. E. N., Towers, S. K., Rozov, A., Buhl, E. H., Whittington, M. A., & Monyer, H. (2001). Impaired electrical signaling disrupts gamma frequency oscillations in connexin 36-deficient mice. Neuron, 31(3), 487–495. 10.1016/S0896-6273(01)00387-7

Juszczak, G. R., & Swiergiel, A. H. (2009). Properties of gap junction blockers and their behavioural, cognitive and electrophysiological effects: Animal and human studies. Progress in Neuro-Psychopharmacology and Biological Psychiatry, 33(2), 181–198. 10.1016/j.pnpbp.2008.12.014

Landisman, C. E., Long, M. A., Beierlein, M., Deans, M. R., Paul, D. L., & Connors, B. W. (2002). Electrical Synapses in the Thalamic Reticular Nucleus. 22(3), 1002–1009.

Lasseigne, A. M., Echeverry, F. A., Ijaz, S., Michel, J. C., Martin, E. A., Marsh, A. J., Trujillo, E., Marsden, K. C., Pereda, A. E., & Miller, A. C. (2021). Electrical synaptic transmission requires a postsynaptic scaffolding protein. ELife, 10. 10.7554/ELIFE.66898

Lee, E., & Terman, D. (2013). Stable antiphase oscillations in a network of electrically coupled model neurons. In SIAM Journal on Applied Dynamical Systems (Vol. 12, Issue 1, pp. 1– 27). 10.1137/120863083

Lee, S. C., Patrick, S. L., Richardson, K. A., & Connors, B. W. (2014). Two functionally distinct networks of gap junction-coupled inhibitory neurons in the thalamic reticular nucleus. The Journal of Neuroscience : The Official Journal of the Society for Neuroscience, 34(39), 13170–13182. 10.1523/JNEUROSCI.0562-14.2014

Lefler, Y., Yarom, Y., & Uusisaari, M. Y. (2014). Cerebellar inhibitory input to the inferior olive decreases electrical coupling and blocks subthreshold oscillations. Neuron, 81(6), 1389– 1400. 10.1016/j.neuron.2014.02.032

Li, X., Olson, C., Lu, S., Kamasawa, N., Yasumura, T., Rash, J. E., & Nagy, J. I. (2004). Neuronal connexin36 association with zonula occludens-1 protein (ZO-1) in mouse brain and interaction with the first PDZ domain of ZO-1. The European Journal of Neuroscience, 19(8), 2132–2146. 10.1111/J.0953-816X.2004.03283.X

Long, M. A., Jutras, M. J., Connors, B. W., & Burwell, R. D. (2004). Electrical synapses coordinate activity in the suprachiasmatic nucleus. Nature Neuroscience 2004 8:1, 8(1), 61–66. 10.1038/nn1361

Lyons, D. J., Horjales-Araujo, E., & Broberger, C. (2010). Synchronized Network Oscillations in Rat Tuberoinfundibular Dopamine Neurons: Switch to Tonic Discharge by Thyrotropin-Releasing Hormone. Neuron, 65(2), 217–229. 10.1016/j.neuron.2009.12.024

Manor, Y., Rinzel, J., Segev, I., & Yarom, Y. (1997). Low-amplitude oscillations in the inferior olive: a model based on electrical coupling of neurons with heterogeneous channel densities. Journal of Neurophysiology, 77(5), 2736–2752. 10.1152/JN.1997.77.5.2736

Miller, A. C., & Pereda, A. E. (2017). The electrical synapse: Molecular complexities at the gap and beyond. Developmental Neurobiology, 77(5), 562–574. 10.1002/DNEU.22484;ISSUE:ISSUE:DOI

MV, B. (1997). Gap junctions as electrical synapses. Journal of Neurocytology, 26(6), 349– 366. 10.1023/A:1018560803261

Nagy, J. I., Pereda, A. E., & Rash, J. E. (2018). Electrical synapses in mammalian CNS: Past eras, present focus and future directions. Biochimica et Biophysica Acta - Biomembranes, 1860(1), 102–123. 10.1016/J.BBAMEM.2017.05.019

Oku, Y., Hülsmann, S., Zhang, W., & Richter, D. W. (1999). Modulation of glycinergic synaptic current kinetics by octanol in mouse hypoglossal motoneurons. Pflugers Archiv : European Journal of Physiology, 438(5), 656–664. 10.1007/S004249900089

Placantonakis, D. G., Bukovsky, A. A., Aicher, S. A., Kiem, H. P., & Welsh, J. P. (2006). Continuous electrical oscillations emerge from a coupled network: a study of the inferior olive using lentiviral knockdown of connexin36. The Journal of Neuroscience : The Official Journal of the Society for Neuroscience, 26(19), 5008–5016. 10.1523/JNEUROSCI.0146-06.2006

Qi-Lytle, X., Sayers, S., & Wagner, E. J. (2023). Current Review of the Function and Regulation of Tuberoinfundibular Dopamine Neurons. International Journal of Molecular Sciences 2024, Vol. 25, Page 110, 25(1), 110. 10.3390/IJMS25010110

Ransey, E., Thomas, G. E., Wisdom, E. M., Almoril-Porras, A., Bowman, R., Adamson, E., Walder-Christensen, K. K., White, J. A., Hughes, D. N., Schwennesen, H., Ferguson, C., Tye, K. M., Mague, S. D., Niu, L., Wang, Z. W., Colón-Ramos, D., Hultman, R., Bursac, N., & Dzirasa, K. (2026). Long-term editing of brain circuits using an engineered electrical synapse. In Nature (Vol. 655, Issue July). Springer US. 10.1038/s41586-026-10501-y

Sharp, A. A., Abbott, L. F., & Marder, E. (1992). Artificial electrical synapses in oscillatory networks. Journal of Neurophysiology, 67(6), 1691–1694. 10.1152/jn.1992.67.6.1691

Sherman, A., & Rinzel, J. (1992). Rhythmogenic effects of weak electrotonic coupling in neuronal models. Proceedings of the National Academy of Sciences of the United States of America, 89(6), 2471–2474. 10.1073/pnas.89.6.2471

Söhl, G., Maxeiner, S., & Willecke, K. (2005). Expression and functions of neuronal gap junctions. Nature Reviews Neuroscience, 6(3), 191–200. 10.1038/nrn1627

Srinivas, M., Rozental, R., Kojima, T., Dermietzel, R., Mehler, M., Condorelli, D. F., Kessler, J. A., & Spray, D. C. (1999). Functional Properties of Channels Formed by the Neuronal Gap Junction Protein Connexin36. 19(22), 9848–9855.

Stagkourakis, S., Dunevall, J., Taleat, Z., Ewing, A. G., & Broberger, C. (2019). Dopamine release dynamics in the tuberoinfundibular dopamine system. Journal of Neuroscience, 39(21), 4009–4022. 10.1523/JNEUROSCI.2339-18.2019

Stagkourakis, S., Pérez, C. T., Hellysaz, A., Ammari, R., & Broberger, C. (2018). Network oscillation rules imposed by species-specific electrical coupling. ELife, 7, 1–18. 10.7554/eLife.33144.001

Stagkourakis, S., Smiley, K. O., Williams, P., Kakadellis, S., Ziegler, K., Bakker, J., Brown, R. S. E., Harkany, T., Grattan, D. R., & Broberger, C. (2020). A Neuro-hormonal Circuit for Paternal Behavior Controlled by a Hypothalamic Network Oscillation. Cell, 1–16. 10.1016/j.cell.2020.07.007

Terman, D., Lee, E., Rinzel, J., & Bem, T. (2011). Stability of anti-phase and in-phase locking by electrical coupling but not fast inhibition alone. SIAM Journal on Applied Dynamical Systems, 10(3), 1127–1153. 10.1137/100813774

Vogelsy, T. P., Froemkey, R. C., Doyon, N., Gilson, M., Haas, J. S., Liu, R., Maffei, A., Miller, P., Wierenga, C. J., Woodin, M., Zenke, F., & Sprekelery, H. (2013). Inhibitory synaptic plasticity: Spike timing-dependence and putative network function. *Frontiers in Neural Circuits*, JUNE. 10.3389/FNCIR.2013.00119

Watanabe, A. (1958). The Interaction of Electrical Activity Among Neurons of Lobster Cardiac Ganglion. The Japanese Journal of Physiology, 8, 305–318. 10.2170/jjphysiol.8.305

Zsiros, V., & Maccaferri, G. (2005). Electrical Coupling between Interneurons with Different Excitable Properties in the Stratum Lacunosum-Moleculare of the Juvenile CA1 Rat Hippocampus. 25(38), 8686–8695. 10.1523/JNEUROSCI.2810-05.2005

